# The Modulation of Pain by Circadian and Sleep-Dependent Processes: A Review of the Experimental Evidence

**DOI:** 10.1101/098269

**Authors:** Megan Hastings Hagenauer, Jennifer A. Crodelle, Sofia H. Piltz, Natalia Toporikova, Paige Ferguson, Victoria Booth

## Abstract

This proceedings paper is the first in a series of three papers developing mathematical models for the complex relationship between pain and the sleep-wake cycle. Here, we briefly review what is known about the relationship between pain and the sleep-wake cycle in humans and laboratory rodents in an effort to identify constraints for the models. While it is well accepted that sleep behavior is regulated by a daily (circadian) timekeeping system and homeostatic sleep drive, the joint modulation of these two primary biological processes on pain sensitivity has not been considered. Under experimental conditions, pain sensitivity varies across the 24 h day, with highest sensitivity occurring during the evening in humans. Pain sensitivity is also modulated by sleep behavior, with pain sensitivity increasing in response to the build up of homeostatic sleep pressure following sleep deprivation or sleep disruption. To explore the interaction between these two biological processes using modeling, we first compare the magnitude of their effects across a variety of experimental pain studies in humans. To do this comparison, we normalize the results from experimental pain studies relative to the range of physiologically-meaningful stimulation levels. Following this normalization, we find that the estimated impact of the daily rhythm and of sleep deprivation on experimental pain measurements is surprisingly consistent across different pain modalities. We also review evidence documenting the impact of circadian rhythms and sleep deprivation on the neural circuitry in the spinal cord underlying pain sensation. The characterization of sleep-dependent and circadian influences on pain sensitivity in this review paper is used to develop and constrain the mathematical models introduced in the two companion articles.

## 1 Introduction: A vicious cycle

The experience of pain has a complex relationship with the sleep-wake cycle. Pain serves two important purposes: to motivate individuals to escape and avoid physical insult and to aid in healing by promoting the protection and immobilization of injured body parts. This first purpose necessitates rapid response and arousal, two processes that are suppressed by sleep, whereas the second purpose is closely tied to the concept of rest. Thus pain makes us tired (promotes the homeostatic drive to sleep), and increased sensitivity to pain during the night is coordinated with our daily circadian rhythm to promote immobilization and healing during the rest period [3]. However, the presence of pain is arousing and can inhibit our ability to initiate and maintain sleep, especially the deeper recuperative stages of sleep [23]. When sleep is disrupted or limited, the perception of pain further intensifies, healing is delayed, and pathological processes promoting the development of chronic pain can proceed unchecked [12]. Within clinical settings, this progression of events can create a vicious cycle of inadequate pain management [23], which is further complicated by similarly strong interdependencies between the sleep-wake cycle and the effectiveness of most forms of analgesia [3, 12, 23].

The development and analysis of mathematical models of this vicious cycle can lead to better understanding of the interactions between sleep and pain, which could improve pain management. In this article, we review the experimental and clinical evidence documenting the modulation of pain by sleep and circadian processes in humans and animals and introduce a novel analysis of this data that is used to justify and constrain the mathematical models introduced in the companion articles.

## 2 What is pain?

“Pain is an unpleasant sensory and emotional experience associated with actual or potential tissue damage, or described in terms of such damage" according to the International Association for the Study of Pain [26]. Pain can be caused by different types of actual or potential tissue damage, including adverse temperature conditions (heat, cold), intense mechanical stimulation or pressure, electric shock, constricted vasculature, or chemical irritation, as well as processes generated within the body, such as inflammation and pathological nerve damage (neuropathy). Pain can be derived experimentally or from natural conditions, and can occur on a variety of time scales. Experimental studies of “acute” pain sensitivity typically induce brief (“phasic”), localized, superficial pain to peripheral tissues. Such brief stimulation actually consists of two sensations: a fast, sharp pain and a slower, dull pain. Occasionally, experimental studies will induce longer duration (“tonic”) acute pain that can last for hours [24]. Within clinical settings, chronic pain conditions can last for months or years.

As there are different types of pain that can be felt, there are different ways in which the body receives and processes pain signals. Sensory neurons (afferent neurons) in the peripheral nervous system sense stimuli and send that information to the spinal cord for processing. These neurons and their nerve fibers are specialized for detecting innocuous or noxious stimuli. Non-painful touch sensations are transmitted by A*β* afferent fibers while there are two major classes of nociceptive (pain-receptive) afferent fibers: A*δ* and C. Medium diameter A*δ* fibers mediate localized, sharp, fast pain sensations, while small diameter C fibers mediate the more diffuse and duller slow pain sensations [11]. The “fast pain” A*δ* fibers are wrapped in a fatty sheath called myelin that allows for rapid transmission of signals, at speeds of 4 to 30 m/sec. This is also true for the A*β* fibers. In contrast, the “slow pain” C fibers are not myelinated and, due to their small diameter, transmit signals at speeds of less than 2 m/sec [24].

Different types of nerve fibers report to different areas in the spinal cord. In general, sensory neurons have their cell bodies in the dorsal root ganglia, a cluster of nerve cell bodies located in the spinal cord. Primary afferent fibers morphologically differ from other nerve fibers in that their axons and dendrites, usually responsible for sending and receiving signals, respectively, have equivalent biochemical makeup and thus these neurons can send and receive signals through both their axons and dendrites [1]. Signals in these afferent fibers are transmitted to the dorsal horn of the spinal cord, an area that is responsible for receiving information from the sensory neurons, processing it, and sending signals up to the brain. The dorsal horn contains many populations of neurons, including excitatory and inhibitory interneurons. One such population of neurons in the dorsal horn, called the Wide Dynamic Range (WDR) neurons, receive direct inputs from the touch and nociceptive afferent fibers as well as inputs from interneuron populations, and constitute the primary output from the dorsal horn to the brain. As such, pain intensity is correlated with the firing rate and the duration of firing of the WDR neurons.

Since pain is both an unpleasant sensory and emotional experience, pain-related input from the spinal cord engages multiple neural circuits in the brain, including the brainstem, thalamus, and cortex. These circuits involve a wide range of neurotransmitter systems, including the well-studied opioid system. Many of these higher-level cognitive and emotional responses to pain exert their influence over pain perception via descending projections to the dorsal horn of the spinal cord. This “top-down” feedback on sensory processing can act to either inhibit or facilitate pain sensation, essentially providing a “gate” for the transmission of nociceptive information to the brain [25]. Thus, there is a tradition of modeling pain processing by focusing exclusively on spinal cord circuitry.

## 3 The relationship between the sleep cycle and pain sensitivity in humans

The daily timing of sleep is widely accepted as an interaction between two independent processes: a homeostatic drive to sleep, which builds up over the course of wakefulness in a saturating manner and dissipates during sleep, and a circadian timing system, which rhythmically influences the levels of sleep drive required to initiate and maintain sleep [8]. When exploring the literature documenting the relationship between sleep and pain, we found that the influences of both circadian rhythms and homeostatic sleep drive were rarely measured within the same experiment, despite ample evidence that both processes modulate pain sensitivity [3, 12, 23]. Instead, experimental studies tended to fall into two broad categories. In one variety of experiment, pain perception was measured across the day (24 hours) in subjects maintaining their normal sleep schedule. Therefore, the data in these experiments should represent a combination of the influences of time-of-day and a normal modest 16-hour build-up of homeostatic sleep drive during waking and 8 hour dissipation of homeostatic sleep drive during sleep. In the other variety of experiment, subjects were sleep deprived for 1-3 nights or had their sleep restricted to less than a typical 8 hours, and pain perception was recorded at various times. In these experiments, there should be a large build up of homeostatic sleep drive, the effects of which may be more or less obvious at different times of day due to circadian modulation. We review these two forms of data below and introduce a novel analysis of the data that allows a comparison of results from these two categories of experiments and from studies using different pain modalities. For the sake of simplicity, we focus primarily on data derived from studies using pain modalities of experimentally-induced brief (acute/phasic), superficial pain to peripheral tissues.

### 3.1 There is a daily rhythm in experimental pain sensitivity in humans

Pain sensitivity follows a daily cycle in many clinical conditions [3], but it is currently unclear how much of that rhythmicity is derived from daily fluctuation in the underlying causes driving the pain (for example, nocturnal release of oxytocin induces contractions during labor) versus rhythmicity in the neural processing of pain. Within the experimental pain literature, rhythmic influences on pain sensation occur regardless of whether pain responses are measured subjectively or objectively [7, 2, 9, 36], suggesting that the rhythmic modulation of pain responses occurs at a basic physiological level. This rhythmic modulation of pain sensitivity increases with pain intensity [15,20,9], so that the more intense the pain is overall, the greater the change in the person’s sensitivity to the pain across the day. Rhythmic influences on pain sensitivity are detectable in experiments involving a variety of different kinds of painful stimuli, including cold, heat, current, pressure, and ischemia (Table 1, Table 2). These stimuli are found to be most painful during hours of the day when experimental subjects are likely to be tired – late afternoon, evening, and night (Table 1).

**Table 1:**
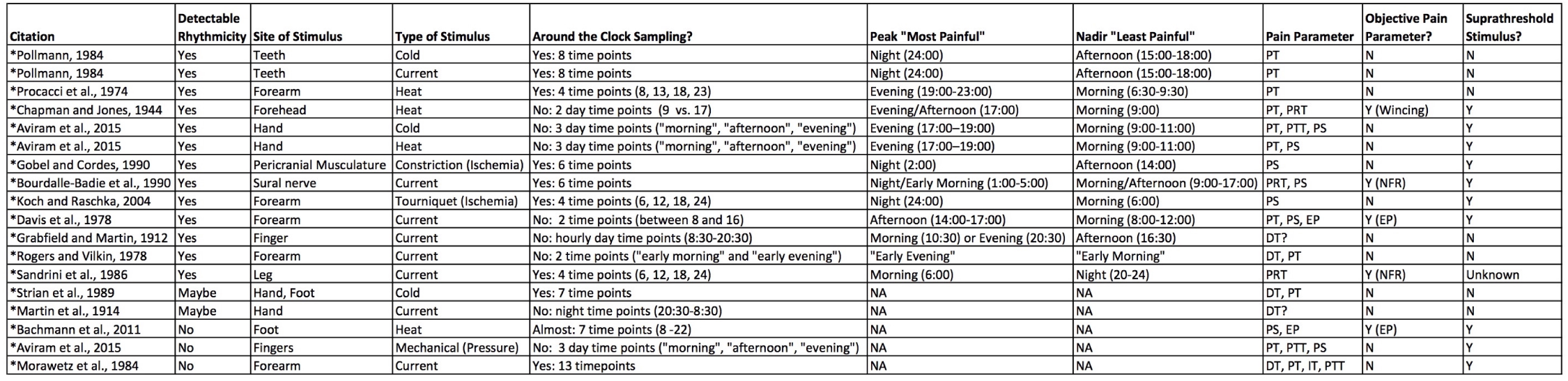
Studies measuring daily rhythms in experimental human pain sensitivity. Note that these studies focus on brief (acute, phasic) superficial pain in peripheral tissues. Clock hours are in military time (0:00-24:00). Abbreviations: DT: Detection threshold, PT: Pain threshold, PRT: Pair response threshold, PS: Pain sensitivity (intensity ratings), PTT: Pain tolerance threshold, IT: Intervention threshold, NFR: Nociceptive flexion reflex, EP: Somatosensory evoked potential (EEG).

**Table 2:**
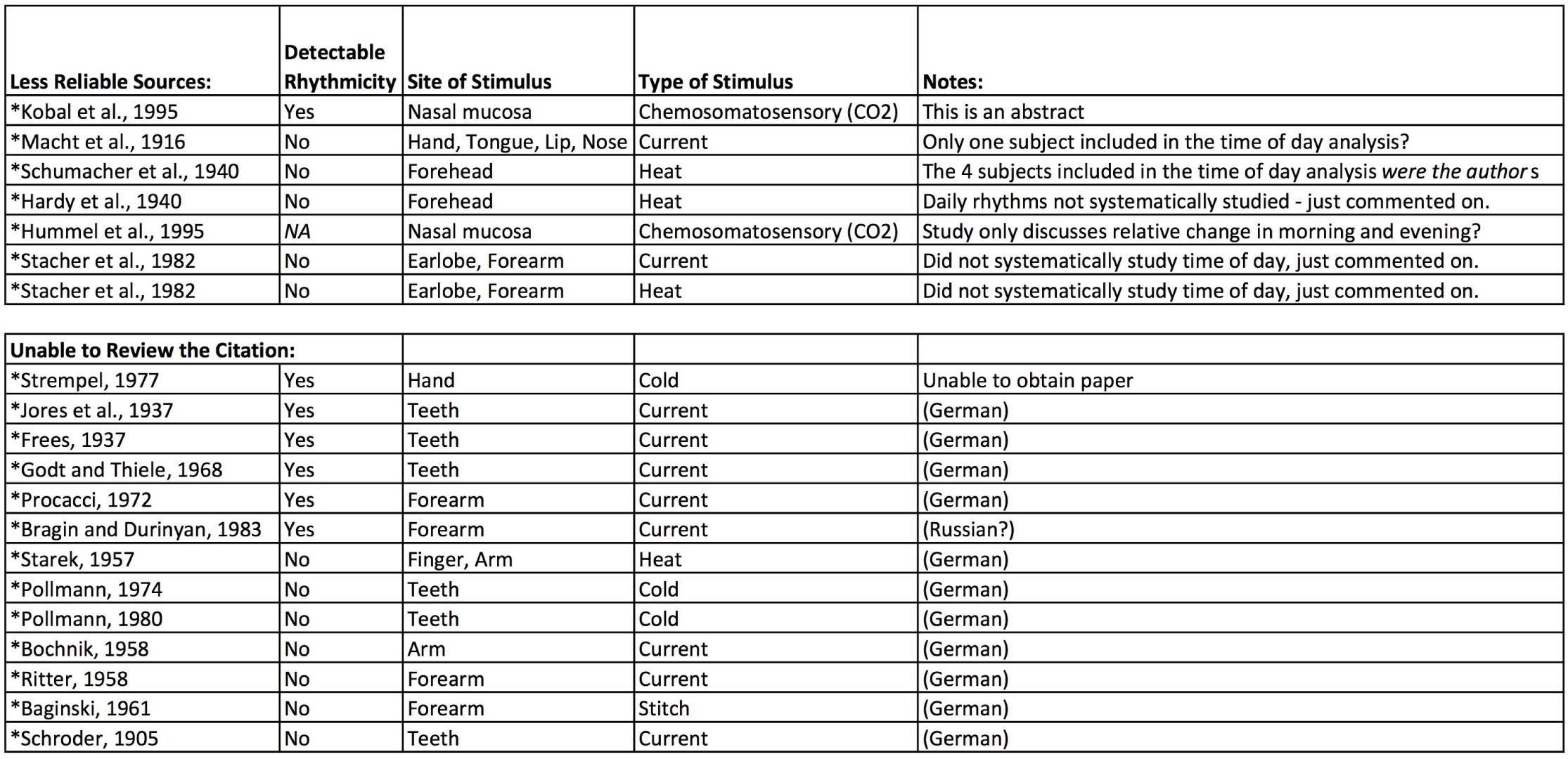
Studies measuring daily rhythms in experimental human pain sensitivity that have been reviewed in previous publications [15, 3] but that we were either personally unable to review or found to be unreliable.

To better characterize this rhythm, we constructed a prototypical “daily pain sensitivity” function by drawing data from four high-quality experiments that measured pain sensitivity at multiple time points around the 24-hr day using diverse testing procedures:

1. The threshold for nociceptive pain reflex in response to electrical current (n=5, [2]), an objective measure of pain sensitivity. In this study, measurements were taken from the same subjects every 4 hours within a consecutive 24 hour laboratory study (beginning at 13:00). The study states that subjects lived in “elementary conditions of social synchronization (08:00-23:00)” and remained in bed during night measurements.
2. The threshold for tooth pain in response to cold (n=79, [31]), and the threshold for tooth pain in response to electrical stimulation (n=56, [31]). In this large study, measurements were taken from the same subjects every 3 hrs across a 24 hr day. From the methods, it is unclear whether these measurements were completed consecutively, but in a follow-up study in the same paper using a smaller sample size they replicate their results using measurements taken at 24+ hr intervals. During the tests, the subjects maintained their normal living cycles.
3. The threshold for forearm pain in response to heat (n=39, [32]). In this large study, measurements were taken from the same subjects at 4 time points across a 24 hr day (8:00, 13:00, 18:00, 23:00). In women, this procedure was repeated at 3 different points across their menstrual cycle (days 7, 15, and 23). During the experiment, subjects maintained their normal daily routine (sleeping hours 24:00-7:00).

For each study, we only had access to the summary data presented in the figures. Using these data, we standardized the pain measurements by converting them to percent of mean (or the mesor of the depicted rhythm). For ease of use, we inverted measures of pain threshold to pain sensitivity so that low pain threshold corresponded to high pain sensitivity. Thus, in our function, high measurement values are associated with greater pain. Time was standardized in relation to either scheduled or estimated morning wake time in order to organize the data in a manner more akin to the “zeitgeber time” used in sleep and circadian literature.

Following these transformations, the data collectively formed a tight curve that resembled a sinusoid. To produce a smoothed version of the curve for later use as our model input, we used the loess function in R 3.2.1 (loess{stats}), R Core Team 2014), which is a form of local polynomial regression that resembles a “vertical sliding window that moves across the horizontal scale axis of the scatterplot” [18].The benefit of using loess() is that it does not assume a functional form for the relationship between X and Y and therefore, to some degree, “allows the data to speak for themselves”[18]. A traditional equation with coefficients is not produced. There is a parameter (alpha, sometimes called span) that controls the degree of smoothing via the width of the sliding window. The larger the alpha value, the smoother the curve. If alpha is too small, overfitting is possible.We used the default (alpha=0.75). There is also a parameter (lambda) that specifies the degree of the polynomial. We used lambda = 2, meaning that quadratic equations were used, which can better capture “peaks” and “valleys” (local minima/maxima) in the data [18]. Using this unbiased approach, we still found that the output curve strongly resembled a simple sinusoid (R2loess=0.64, Figure 1).

**Fig. 1:**
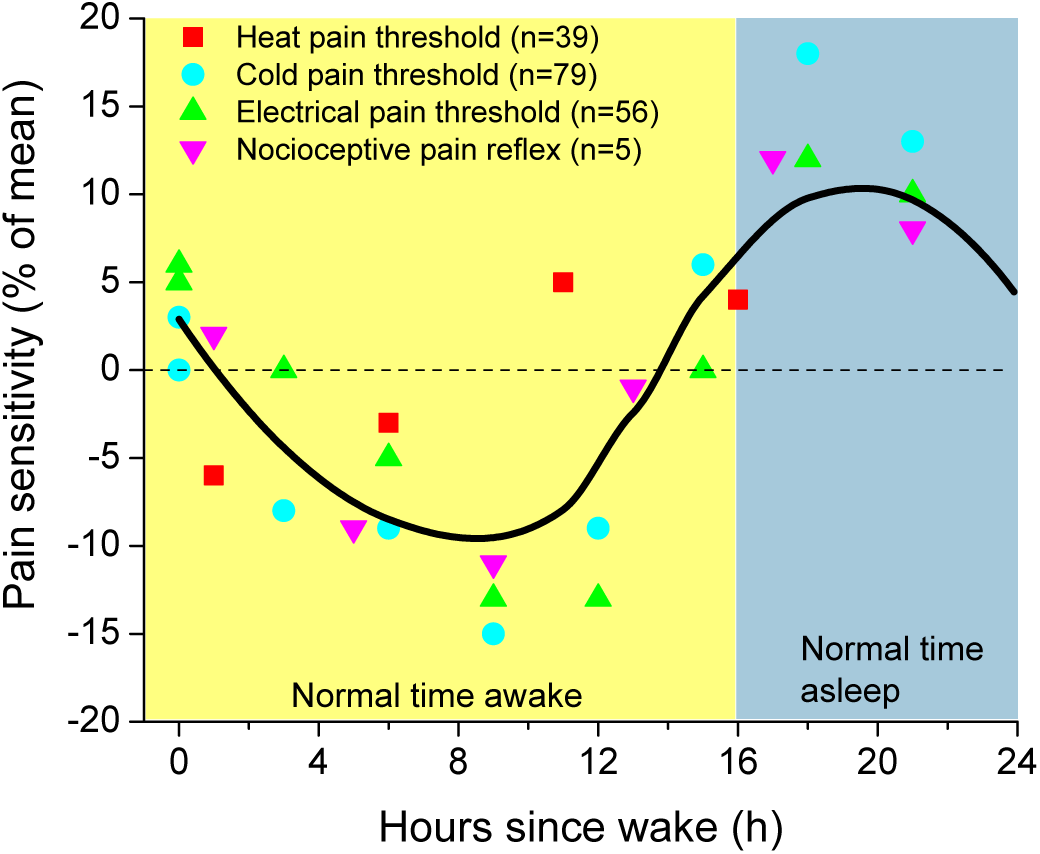
Prototypical human “daily pain sensitivity” curve constructed from summarized data from four high-quality experimental studies of pain responses [2,31, 32]. Time was standardized in relation to either scheduled or estimated morning wake time (time of wake = 0). Each point represents the mean value for that time point as derived from the published figures in each study, converted to a percent of the study’s overall mean (or the mesor of the depicted rhythm). For ease of use, we inverted measures of pain threshold, so that low pain thresholds are presented in the graph as high “pain sensitivity”. The smoothed curve was produced using an unbiased loess{stat} regression in R.

We hypothesize that this best-fit curve represents an average daily rhythm in pain sensitivity for humans and that the rhythm is affected by both homeostatic sleep drive and a circadian rhythm in pain sensitivity. The curve has a sharp peak in pain sensitivity occurring close to sleep onset (18 hours following wake, or approximately 1am) and then decreases during the night. This is consistent with an effect of homeostatic sleep drive on pain sensitivity, and fits previous demonstrations that mental fatigue can decrease pain threshold (i.e., increase pain sensitivity) by 8 – 10% [7]. The curve also has a distinct trough in pain sensitivity in the afternoon (following 9 hours of wake, or approximately 4pm). This pattern does not fit what would be expected due to an effect of homeostatic sleep drive and instead suggests the influence of a circadian rhythm.

### 3.2 Homeostatic sleep drive increases pain sensitivity in humans

Within the clinical literature, there are at least 14 studies demonstrating that the experience and intensity of pain correlate with sleep duration or quality [12]. However, the causal nature of this relationship is best evaluated within controlled experiments, and these experimental results have been wide-ranging. Within human experiments, sleep deprivation or restriction produced no effect on experimental pain [10, 39], small 2 – 10% increases in pain [29, 21, 37], or much larger 18 – 118% increases in pain [43, 34, 38]. The diversity of these effects may be due to the variety of sleep protocols used (as suggested by [38]) or the cognitive and emotional context accompanying each experiment (e.g., [39]). Even within a particular protocol, the intensity or quality of experimental pain may determine the impact of sleep deprivation, with one study observing increases in experimental pain that ranged from 6 – 118% depending on the method used to inflict and measure pain [38] (Table 3).

**Table 3:** The estimated impact of sleep deprivation on experimental pain can vary greatly across experimental pain measures within the same study [38]. In this study, the responses of 14 healthy subjects to five different measurements of evoked pain were assessed after a night of undisturbed sleep (control) and after a night of total sleep deprivation (Total Sleep Dep). The change in response is measured as a percentage of the control response.

| Schuh-Hofer et al., 2013 | Control | Total Sleep Dep | Difference | % Increase in Pain |
| --- | --- | --- | --- | --- |
| Cold Pain Threshold (degrees C) | 14.7 | 20.4 | 5.7 | 38.8 |
| Heat Pain Threshold (degrees C) | 44.2 | 41.4 | 2.8 | 6.3 |
| Pressure Pain Threshold (kPa) | 446.6 | 376.6 | 70 | 18.6 |
| Mechanical Pain Threshold (mN) | 67.8 | 43.5 | 24.3 | 55.9 |
| Mechanical Pain Sensitivity (NRS 0-100) | 5.5 | 12 | 6.5 | 118.2 |

However, we noted that much of the variability in pain sensitivity across studies could be accounted for by the method of normalization used to compare data. For example, when using percentage change as our standardized unit, we can artificially see a larger effect of sleep deprivation on cold pain threshold if the original units are in degrees Celsius instead of in degrees Fahrenheit (Table 4).

**Table 4:** An example of why it is difficult to compare magnitude of effect across different units for measuring pain [38]. Changes in the threshold temperature for evoked pain by cold stimulus to the hand was measured after a night of undisturbed sleep (control) and after a night of total sleep deprivation (Total Sleep Dep). When measured as a percentage of the control response, the same change in response is computed as a larger percentage change when degrees Celsius are used compared to degrees Farenheit.

| Schuh-Hofer et al., 2013 | Control | Total Sleep Dep | % Increase in Pain |
| --- | --- | --- | --- |
| Cold Pain Threshold (degrees C) | 14.7 | 20.4 | 38.8 |
| Cold Pain Threshold (degrees F) | 58.5 | 68.7 | 17.6 |

Similarly, across studies, the effect of sleep deprivation in percentage change units was consistently larger on cold pain threshold than on heat pain threshold, simply because the temperature values for cold pain threshold under the control condition were lower, making the denominator in the percentage change equation (i.e. (change from control threshold temperature) / (control threshold temperature)) smaller. Likewise, percentage change increases in subjective rating scales were almost always exaggerated, since ratings from control subjects were often extremely low, making the denominator in the percentage change equation diminutive.

The typical rationale for using percentage change units for comparing data of different units is the idea that the biological impact of changes in the unit depends on its initial values. For example, if a disease condition increases the number of mRNA transcripts for a particular gene from 100 to 110, this is likely to matter more biologically than an increase from 1000 to 1010. It is not clear that this logic holds true for the units used in pain research (as indicated by the particularly irrational examples above). What is likely to matter more biologically is the percentage of the range of stimulation possible before tissue is genuinely damaged. For example, in a heat threshold experiment, you would expect that pain sensation might reflect a range of temperatures between physiological levels (37 degrees C) and a level of heat that rapidly causes damage (60 degrees C) [51]. In that case, a drop in pain threshold of 3 degrees would cover 13% of the full range of pain sensation possible ((3 degrees)/(60-37 degrees C)=0.13). This is a much more interpretable value than simply saying that a drop in pain threshold from 49 degrees in controls to 46 degrees following sleep deprivation is a 6% change from the original pain threshold ((49-46 degrees)/49 degrees=0.061).

Using this logic, we found that the data documenting the impact of sleep deprivation on experimental pain was much more consistent than it initially appeared. For the different experimental pain measures used in the study of [38], we determined minimum and maximum response values corresponding to the absence of stimulation and the value when tissue damage would occur, respectively. The full range of response values was computed as the difference between the maximum and minimum response values.We then computed the observed difference in response values between the control and total sleep deprivation conditions, as listed in Table 3, as a percentage of the full range of stimulation. Computed in this way, all experimental pain measures indicate that sleep deprivation increases pain by ∼ 5 – 12% of the full range of painful stimulation (Table 5).

**Table 5:** Within a study, the estimated impact of sleep deprivation on experimental pain is quite consistent if the data are normalized as a percentage of the full estimated range for painful sensation in that modality [38]. The observed difference in response values between the control and total sleep deprivation conditions, as listed in Table 3, was computed as a percentage of the full range of stimulation response values.

|  | Observed Difference | Minimum Pain (no stimulation) | Maximum Pain (threshold for tissue damage) | Full Range of Stimulation (Max-Min) | % Range of Stimulation |
| --- | --- | --- | --- | --- | --- |
| Schuh-Hofer et al., 2013 |  |  |  |  |  |
| Cold Pain Threshold (degrees C) | 5.7 | 37 | -51 | 88 | 6.5 |
| Heat Pain Threshold (degrees C) | 2.8 | 37 | 60 | 23 | 12.2 |
| Pressure Pain Threshold (kPA) | 70 | 0 | 686 | 686 | 10.2 |
| Mechanical Pain Threshold (mN) | 24.3 | 0 | 512 | 512 | 4.7 |
| Mechanical Pain Sensitivity (NRS 0-100) | 6.5 | 0 | 100 | 100 | 6.5 |

To apply this logic to multiple studies on the effects of sleep deprivation and circadian modulation of pain, we estimated minimum and maximum stimulation levels necessary to produce a full range of pain responses to a number of different experimental pain modalities, as well as typical pain thresholds (Table 6). From these measurements, we computed the range of physiologically-meaningful stimulation as the difference between the maximum and minimum values, and the range of painful stimulation as the difference between the maximum and pain threshold values. We then normalized the results across studies by converting changes in pain response values to percentage changes within the full range of physiologically-meaningful stimulation or percentage changes within the range of painful stimulation. Following this normalization, the magnitudes of the effects of sleep deprivation and daily rhythms were less variable across studies. This implied that normalizing data based on percentage changes within the range of painful stimulation was superior to using a simple percentage of the mean. (However, please note that the data necessary to perform this improved normalization were not available for all studies - for example, several studies used to construct Figure 1). Using this improved normalization method, we also found that the magnitude of the effects of sleep deprivation and daily rhythms were roughly equivalent (Table 7). Specifically, we found that, on average, evoked pain responses, measured relative to the range of painful stimulation, varied by approximately 14% due to daily rhythms and by approximately 13% in response to sleep deprivation.

**Table 6:** Determining the range of painful stimulation possible within human experimental pain studies before the occurrence of tissue damage, as well as the typical threshold for pain sensation. Sources for computations: *a:* http://www.ehs.neu.edu/laboratory_safety/fact_sheets/-cryogenic_liquids; *b*: [51]; *c*: [41]; *d*: Maximum for Instrument, the typical mA eliciting nociceptive pain reflex by someone who is under general anesthesia for surgery [46]; *e*: Pressure of about 100 lb/in^2^ (7 kg/cm^2^) is required to penetrate the epidermis (1 kg/cm^2^ = 98.07 kPA) [4]; *f*: Maximum for Instrument [38]; *g*: Maximum for Test, rated “Very strong pain" by all participants [15]; *h*: [38]; *i*: [43];*j*: [2];*k*: [20].

| Type of Pain Measurement | Minimum Pain (no stimulation) | Maximum Pain (threshold for tissue damage) | Full Range of Meaningful Stimulation (Max - Min) | Typical Pain Threshold | Pain Threshold as % of Range of Painful Stimulation | Range of Painful Stimulation (Max - Threshold) |
| --- | --- | --- | --- | --- | --- | --- |
| Cold (degrees C)<sup>a</sup> | 37 | -51 | 88 | 12 | 28% | 59 |
| Heat (degrees C)<sup>b</sup> | 37 | 60 | 23 | 46 | 37% | 15 |
| Heat (mcal/cm<sup>2</sup>)<sup>c</sup> | 0 | 2420 | 2420 | 871 | 36% | 1540 |
| Electric Stimulation of Sural Nerve (mA)<sup>d</sup> | 0 | 40 | 40 | 12 | 30% | 28 |
| Pressure (kPa)<sup>e</sup> | 0 | 686 | 686 | 447 | 65% | 240 |
| Mechanical (mN)<sup>f</sup> | 0 | 512 | 512 | 68 | 13% | 444 |
| Tolerable Pressure Duration (sec)<sup>g</sup> | 0 | 360 | 360 | 15 | 13% | 345 |
| Mechanical Pain Sensitivity (NRS 0-100)<sup>h</sup> | 0 | 100 | 100 | 1 | 1% | 99 |
| Laser Heat VAS Pain Ratings (0-100)<sup>i</sup> | 0 | 100 | 100 | 1 | 1% | 99 |
| Electrical Stimulation Pain Ratings (0-10)<sup>j</sup> | 0 | 10 | 10 | 1 | 10% | 9 |
| Tourniquet Pain Intensity Ratings (0-10)<sup>k</sup> | 0 | 10 | 10 | 1 | 10% | 9 |

**Table 7:** Experimental results illustrating the effects of daily rhythm and sleep deprivation on pain thresholds in humans for multiple pain modalities. These results are normalized with respect to the range of physiologically-meaningful or painful stimulation computed in Table 6. Across studies, the estimated impact of the daily rhythm and sleep deprivation on experimental pain is quite consistent if the data are normalized as a percentage of either the estimated range for physiologically-meaningful sensation or painful sensation in that modality.

| Type of Pain Measurement: | In Original Units: |  | As % of Mean: |  | As % of Full Range (max-min): |  | As % of Painful Range (max-threshold): |  |
| --- | --- | --- | --- | --- | --- | --- | --- | --- |
|  | Daily Rhythm: | Sleep Deprivation: | Daily Rhythm: | Sleep Deprivation: | Daily Rhythm: | Sleep Deprivation: | Daily Rhythm: | Sleep Deprivation: |
| Cold (degrees C) [38, 21] |  | 3.7 |  | 32% |  | 4% |  | 6% |
| Heat (degrees C) [38, 21, 37] |  | 2.8 |  | 6% |  | 12% |  | 19% |
| Heat (mcal/cm<sup>2</sup>) [32] | 122 |  | 14% |  | 5% |  | 8% |  |
| Electric Stimulation of Sural Nerve (mA) [2] | 2.4 |  | 20% |  | 6% |  | 9% |  |
| Pressure (kPA) [38] |  | 70.0 |  | 16% |  | 10% |  | 29% |
| Mechanical (mN) [38] |  | 24.3 |  | 36% |  | 5% |  | 5% |
| Tolerable Pressure Duration (sec) [15] | 41 |  | 36% |  | 11% |  | 12% |  |
| Mechanical Pain Sensitivity (NRS 0-100) [38] |  | 6.5 |  | 118% |  | 7% |  | 7% |
| Laser Heat VAS Pain Ratings (0-100) [43] |  | 9.5 |  | 23% |  | 10% |  | 10% |
| Electrical Stimulation Pain Ratings (0-10) [2] | 2.3 |  | 48% |  | 23% |  | 26% |  |
| Tourniquet Pain Intensity Ratings (0-10) [20] | 1.45 |  | 18% |  | 15% |  | 16% |  |
|  | AVERAGE |  | 27% | 38% | 12% | 8% | 14% | 13% |
|  | STDEV |  | 14% | 40% | 7% | 3% | 7% | 10% |
|  | Signal/Noise |  | 1.92 | 0.95 | 1.65 | 2.45 | 1.94 | 1.33 |

### 3.3 A cross-species comparison: circadian rhythms and homeostatic sleep drive influence pain sensitivity in laboratory rodents

The vast majority of what is known regarding the influence of circadian rhythms and sleep-wake cycles on pain processing circuitry comes from studies on laboratory rodents. In order to properly compare these data with that of humans, it is important to understand that the circadian and sleep systems of laboratory rodents differ from humans in several fundamental ways. To begin with, laboratory mice and rats are nocturnal, which means that most of their wakefulness occurs at night and most of their sleep occurs during the day. They are also polyphasic sleepers, which means that they sleep in short, multi-minute bouts, interrupted by waking, and rarely exhibit consolidated wakefulness that extends beyond several hours. Despite their unconsolidated wake and sleep, they still generally exhibit a progressive build-up of homeostatic sleep drive across the nighttime active period, and dissipation during the daytime rest period [44].

Similar to humans, there is clear evidence that pain sensation in laboratory rodents is modulated by both time-of-day [28, 6, 35, 52, 13, 33, 22, 42, 19] and sleep deprivation [17, 49, 48, 47, 45, 27]. Unlike humans, we can easily place laboratory rodents into constant environmental conditions and thus be able to demonstrate with certainty that the influence of time-of-day on pain sensation is due to an endogenous circadian clock instead of simple passive responses to a rhythmic environment [33]. However, the timing of the daily peak in pain sensitivity varies in different strains of inbred rodents by as much as 12 hours [6], making it sometimes difficult to draw generalized conclusions about the influence of circadian rhythms on pain sensitivity. Another notable difference between humans and rodents is that the duration of sleep deprivation necessary to observe an effect on pain responses is much smaller, since rodents typically do not exhibit consolidated wakefulness on the scale of multiple hours.

## 4 Circadian rhythms and homeostatic sleep drive modulate pain neural circuitry

The neural location for the circadian modulation of pain begins at the most fundamental level of the pain circuitry: sensory afferent input into the spinal cord. Within the dorsal root ganglia, which are the neural structures that contain the cell bodies for the sensory afferent neurons, there is clear evidence for endogenous circadian rhythmicity. The dorsal root ganglia rhythmically express a full complement of clock genes, which are the genes responsible for generating daily rhythmicity throughout the body [52]. The dorsal root ganglia also demonstrate rhythmic expression of genes necessary for synaptic transmission, including voltage-gated calcium channel subunits [22] and NMDA glutamate receptor subunits [52]. However, since the dorsal root ganglia contain the cell bodies for a wide variety of afferent neurons, it could be argued that measuring rhythmicity in the dorsal root ganglia as a whole does not necessarily indicate that the nociceptors responsible for pain transmission are rhythmic. Two pieces of evidence suggest otherwise. First, an estimated 82% of afferents are nociceptors [11], thus it is likely that the majority of mRNA collected from the dorsal root ganglia in these experiments represents mRNA from pain transmitting cells. Second, researchers have discovered rhythmic expression of the mRNA and protein for Substance P, a neurotransmitter important for pain-signaling from C fibers [52]. Therefore, it is likely that the nociceptive afferent neurons themselves are rhythmic. That said, the cell bodies for the non-nociceptive fibers in the dorsal root ganglia probably also contain endogenous rhythmicity. In human studies the influence of time-of-day on non-noxious mechanical sensitivity, which is conveyed by A*β* fibers, differs from that of painful stimuli, which is conveyed by A*δ* and C fibers, with the rhythm in mechanical sensitivity peaking in the late afternoon (15:00-18:00) and the rhythm in pain sensitivity peaking in the middle of the night (between midnight and 03:00 [31]).

The top-down inhibition of pain processing in the dorsal horn also exhibits a daily rhythm. In humans, placebos best alleviate pain in the early afternoon [31]. In laboratory rodents, there is a daily rhythm in the strength of stress-induced analgesia and endogenous opioid-release, and this rhythm persists under constant environmental conditions [33, 50]. Opioid receptors in the brainstem, which are important for analgesia, exhibit a strong daily rhythm [42]. However, it is possible that these daily rhythms in the top-down inhibition of pain do not represent direct influences of the circadian clock, but instead are a response to the rhythmic build-up and dissipation of homeostatic sleep drive across the day. In support of this theory, there is strong evidence demonstrating that sleep deprivation influences the highest levels of pain processing. Sleep deprivation is already well known to disproportionately affect the energy and resource-needy cortex. Therefore, it is unsurprising that sleep deprivation in humans eliminates distraction-based analgesia [43] and decreases central pain modulation [16, 5]. Even top-down pain inhibition originating from lower levels of the central nervous system is crippled by sleep deprivation, including diffuse noxious inhibitory controls in humans [39, 16] and stress-induced hyperalgesia in rodents [45]. Pharmacological manipulations that mimic top-down pain inhibition, such as morphine, are ineffective following severe sleep deprivation [45, 27].

Sleep deprivation can also alter more fundamental levels of pain processing in the spinal cord, including neurotransmission via glutamate (mGLUR5, NMDA), GABA, and NOS, as well as the passive spread of electric potential via astrocytic gap junctions and the production of reactive oxygen species [49, 48, 47]. Despite these effects, under conditions in which the top-down inhibition of pain is minimal, there seems to be less evidence that homeostatic sleep drive influences pain processing. For example, there is some evidence that sleep deprivation may not affect the processing of fast pain (*Aδ* input). Cortical responses to fast pain actually diminish following sleep restriction [43], and, for faster reflexive behaviors (such as tail withdrawal latency), sleep deprivation effects are sometimes not found [49]. Likewise, homeostatic sleep drive does not seem to contribute much to the typical daily rhythm in acute pain sensitivity in rodents. For example, experiments performed in mice with a critical mutation in the essential clock gene Per2 show a complete elimination of daily rhythms in acute pain under typical housing conditions [30, 52], despite maintaining elevated nocturnal activity in response to the laboratory light-dark cycle [40], and thus presumably also retaining a daily rhythm in the build-up and dissipation of homeostatic sleep drive.

In the case of more severe or chronic pain, the influences of homeostatic sleep drive on top-down inhibition may be more relevant. For example, Per2 mutant mice continued to exhibit daily rhythms in inflammatory pain in a manner that matched a predicted build-up and dissipation of homeostatic sleep drive in response to nocturnal behavioral patterns [52]. Both inflammatory and neuropathic conditions are also characterized by a 8– 12 hour shift in the phasing of daily rhythms in pain sensitivity [52, 42, 22, 14], which may represent an increased influence of homeostatic drive on the top-down inhibition of pain when pain is extended over a longer time scale.

## 5 Discussion

In summary, there is a substantial body of work documenting the effects of both daily rhythms and sleep deprivation on acute pain sensitivity under experimental conditions in humans and rodents. These results appear divergent at first glance, but upon closer inspection seem to generally agree that peak acute/phasic pain sensitivity in humans occurs during the evening. Our data analysis reveals that the influence of both daily rhythms in pain sensitivity and 24 hours of sleep deprivation typically alter pain sensitivity under experimental conditions by 13 – 14% of the full range of painful stimulation. Other studies suggest that the influence of daily rhythms and sleep deprivation may increase with pain intensity. Where these effects originate physiologically is a more recent source of discussion, but it is likely that they represent the intersecting influence of homeostatic sleep drive and the circadian timekeeping system on the central nervous system. There is clear evidence for circadian effects at the level of the spinal cord and there are equally clear effects for sleep-dependent modulation of the top-down inhibition of pain, although it is possible that both processes influence all levels of the central nervous system. Finally, the effect of circadian rhythms and homeostatic sleep pressure on pain sensitivity may differ depending on the type of pain measured, with data clearly indicating that slower C fiber input and tonic pain sensitivity are influenced by both endogenous circadian rhythms and homeostatic sleep drive, whereas fast A*δ* input and faster reflexive pain measures may be less susceptible.

In the companion articles, we introduce two mathematical models to investigate the joint modulation of the circadian rhythm and homeostatic sleep drive on pain. These models address pain at two different levels: at the organismal level as the experience of pain sensitivity and at the neural level as the firing rates of pain processing circuits in the spinal cord.

The organismal-level model addresses a clear gap in our current knowledge: the lack of experimental data measuring the dissociated influence of circadian rhythms and homeostatic sleep drive on pain sensitivity in humans, as would be obtained from a forced desynchrony or constant routine protocol. To address this gap, we adapt the formalism of a classic mathematical model for the regulation of sleep behavior by the circadian rhythm and homeostatic sleep drive, called the Two Process model [8], to simulate the interaction of these two processes on pain sensitivity. The data analysis presented here is used to define a generic “daily pain sensitivity” function (Figure 1), which we decompose into two independent circadian and homeostatic components (Process C and Process S) using a range of potential relative magnitudes constrained to produce results resembling the normalized data in Table 7. Then we use this model to simulate the resultant changes in the daily pain sensitivity rhythm in response to a variety of altered sleep schedules: sleep deprivation, sleep restriction, and shift work.

The neural-level model is based on the circuitry in the dorsal horn of the spinal cord consisting of synaptically coupled populations of excitatory and inhibitory interneurons that process input signals in the primary afferent fibers and influence the output signal of the WDR neurons. The temporal profile of inputs on the different types of afferent fibers and excitability properties of the included neuronal populations are constrained by experimental results. We validate the model by replicating experimentally observed phenomena of A fiber inhibition of pain and wind-up. We then use the model to investigate mechanisms for the observed phase shift in circadian rhythmicity of pain that occurs with neuropathic pain conditions.

In conclusion, while experimental evidence indicates both circadian and sleep-dependent effects on daily pain rhythms, dissecting their interactions that contribute to changes in pain rhythms under varying normal or pathological conditions is difficult experimentally. The mathematical models developed in this series of papers provide frameworks to incorporate the known experimental results of these effects and to investigate their potential interactions under different conditions. By addressing both the behavioral and cellular levels, these models are useful tools to identify how the primary biological processes of sleep, circadian rhythmicity and pain interact.

## 6 Acknowledgements

This work was conducted as a part of A Research Collaboration Workshop for Women in Mathematical Biology at the National Institute for Mathematical and Biological Synthesis, sponsored by the National Science Foundation through NSF Award DBI-1300426, with additional support from The University of Tennessee, Knoxville. This work was additionally partially supported by the following sources: NSF Award DMS-1412119 (VB), NSF RTG grant DMS-1344962 (JC) and the Pritzker Neuropsychiatric Disorders Research Consortium (MH). Any opinions, findings, and conclusions or recommendations expressed in this material are those of the authors and do not necessarily reflect the views of the National Science Foundation.

